# Identification of protective human monoclonal antibodies using a K18 hACE2 transgenic mouse SARS-CoV-2 challenge model

**DOI:** 10.1101/2025.09.13.675985

**Authors:** Bhavesh Borate, Paula A. Pino, Amberlee Hicks, Juan I. Garcia, Andreu Garcia-Vilanova, Chengjin Ye, Jun-Gyu Park, Billie Maingot, Oscar Rodriguez, Tracy Saveria, Drienna Holman, Sharon L. Schendel, Erica Ollman Saphire, Luis Martinez-Sobrido, Jordi B. Torrelles, Andrew Fiore-Gartland

**Author notes:** **Correspondence:** Corresponding Authors: Jordi B. Torrelles and Andrew Fiore-Gartland. Center for Integrated Research in Infectious Diseases (CIRID), Division of Infectious Diseases, Department of Internal Medicine, College of Medicine, The Ohio State University, Columbus, OH, US.

## Abstract

Severe acute respiratory syndrome coronavirus 2 (SARS-CoV-2), the causative agent of coronavirus disease 2019 (COVID-19), uses human angiotensin converting enzyme 2 (hACE2) as its obligate receptor for cell entry. The K18 hACE2 transgenic mouse line, which expresses hACE2 under control of the human keratin 18 (K18) promoter, is used as an animal model for the study of COVID-19 pathogenesis. Here, we evaluate this model in the screening of human monoclonal antibody (hmAb) therapies against SARS-CoV-2. We included 206 hmAbs from the Coronavirus Immunotherapeutic Consortium Database (CoVIC-DB) and identified many that protected against a lethal challenge with the virus. Our data showed that mouse weight change from day 5 onward highly correlated with survival. Many of the protective hmAb candidates we identified also showed strong viral neutralization and spike protein (SP) binding when measured *in vitro*; however, in many cases, *in vitro* assays failed to identify protective hmAbs, suggesting that the mouse model may capture characteristics of the hmAbs that other methods cannot. Our findings demonstrate the relevance of including *in vivo* models for the characterization of therapeutics against SARS-CoV-2, as these improve both accuracy and expediency in the screening process.

## INTRODUCTION

Coronavirus Disease 2019 (COVID-19) is caused by Severe Acute Respiratory Syndrome Coronavirus 2 (SARS-CoV-2), an enveloped RNA virus closely related to SARS-CoV and Middle East Respiratory Coronavirus (MERS-CoV) (1, 2). Like other coronaviruses, SARS-CoV-2 prominently displays a trimeric glycoprotein on its surface, called spike protein (SP), that mediates viral genome entry into host cells via binding and subsequent fusion of the viral membrane with the host cell membrane (2–6). As with SARS-CoV, SARS-CoV-2 uses human angiotensin converting enzyme 2 (hACE2) as its obligate receptor for cell invasion (1, 7, 8). Vaccines and human monoclonal antibodies (hmAb) that target SP to interrupt hACE2 interaction have proven to be effective treatments against COVID-19 (9–12). The prominence of SP on the virion surface and its key role in cell entry render it highly susceptible to immune pressure from the host, leading to rapid evolution of new variants that evade recognition (4, 9, 13–15). This ever-increasing diversity of SARS-CoV-2 propels the need to develop new hmAb therapies to prevent and treat disease as the virus evolves (16). Reliable tools may also be needed to accelerate discovery of Ab therapies when new threats emerge, like novel coronaviruses or other respiratory viruses.

The K18 hACE2 transgenic mouse was developed following the 2003 SARS-CoV outbreak to be used as an animal model for studying SARS-CoV infection in humans (17). Using a C57BL/6 mouse background, the line was engineered to express the hACE2 receptor under the control of the human keratin 18 (K18) promoter for its targeted expression to the airway epithelium (17). Following intranasal (i.n.) inoculation with a human strain of SARS-CoV, mice rapidly developed disease marked by weight loss, lethargy, and death by day 7 post-infection (17). During the early stages of the COVID-19 pandemic, studies showed that following challenge with SARS-CoV-2, these mice succumbed to disease by day 6 post-infection with pathogenesis that partially recapitulated human COVID-19 (18, 19). These and other reports (20, 21) have demonstrated the importance of this mouse model for studying viruses that can use hACE2 for cell entry, which importantly include an array of human and animal coronaviruses with the potential to cause future human pandemics.

The Coronavirus Immunotherapeutic Consortium (CoVIC) is a global partnership that was established during the COVID-19 pandemic to provide a useful compendium of anti-SARS-CoV-2 therapeutic antibodies (22). Candidates submitted to the CoVIC database (CoVIC-DB) were assayed for functional characteristics such as SP binding, hACE2 blockage, and viral neutralization using standardized protocols, allowing for meaningful comparison across multiple potential hmAb therapies. Recently, the Consortium published findings from a multi-lab study comparing anti-SARS-CoV-2 SP candidates submitted to the database for analysis (23). These results showed that *in vitro* SP binding affinity and viral neutralization were both correlated with *in vivo* protection of K18 hACE2 mice against a lethal infection with SARS-CoV-2. Notably, however, potent binding and neutralization *in vitro* did not guarantee protection *in vivo*. Here, we further analyze data from the mouse study to evaluate characteristics of the challenge model for future applications, and we include new data on the binding affinity of mouse serum collected post-treatment to assess inter-mouse variability. Our results confirm that SP binding strength correlates with an antibody’s ability to protect the mice, but we show that *in vitro* assays in general are poor predictors of protection *in vivo*. We also show that by day 5 post-treatment, mouse weight loss is inversely correlated with protection and can provide a more nuanced readout for analyzing antibody effectiveness. These data demonstrate the importance of using both *in vitro* and *in vivo* models for evaluation of Ab-based therapies.

## MATERIALS AND METHODS

### Ethics Statement

Mouse model studies were performed in an animal biosafety level 3 (ABSL3) facility at Texas Biomedical Research Institute (Texas Biomed). Institutional biosafety (IBC) and animal care and use committee (IACUC) protocols for these studies were approved by Texas Biomed under protocols BSC20-004 and IACUC #1745MU in accordance with the National Institute of Health (NIH) and the United States Department of Agriculture (USDA) guidelines.

### Viruses and cells

SARS-CoV-2/human/USA/WA-CDC-WA1/2020 viral strain was obtained from the Biodefense and Emerging Infections Research Resources (BEI Resources). To amplify the virus, Vero-E6 cells sourced from American Type Culture Collection (ATCC) were cultured in Dulbecco’s modified Eagle’s medium (DMEM) plus 10% fetal bovine serum (FBS) at 37°C and 5% CO_2_. Cells were inoculated with virus at a multiplicity of infection (MOI) of 0.001 and cultured for 72 h. Cell culture supernatant was collected and titrated using standard plaque assay. For each challenge experiment, virus was sequenced using next generation sequencing (NGS) and confirmed to be 100% identical to the original BEI Resources stock.

### K18 hACE2 transgenic mouse infection model

For each trial, we used female 6-week-old K18 hACE2 transgenic mice to test 8 different hmAbs (n=10 mice per hmAb) against 3 control groups. Controls for each experiment included infected without treatment (n=5), infected and treated with CC12.3 (positive control previously demonstrated 50% protection at 1.5 mg/mouse, n=10), and a phosphate-buffered saline (PBS)-mock infected and saline treated negative control (n=5). Overall, we obtained validated data from 206 hmAbs. Treatments were provided by intraperitoneal (i.p.) injection with 0.5 mg to 1.5 mg of hmAb obtained from the CoVIC-DB. All hmAbs were unidentified, and experiments were performed in a blinded manner. After 24 h post-treatment, mice were anesthetized and intranasally (i.n.) infected with SARS-CoV-2 virus at a lethal dose of 1×10^5^ PFU/mouse. Peripheral blood was collected just prior to virus challenge to determine hmAb levels by ELISA at the time of infection. All mice were observed for morbidity (body weight loss) and mortality (% survival) signs daily. Mice were humanely euthanized if they presented a loss of 25% of their initial body weight at any time during the study; all remaining mice were humanely euthanized on day 10 post-infection. Blood samples obtained post-treatment and prior to infection were kept at room temperature for 30 min to allow them to clot, followed by centrifugation at 400 x *g* for 15 minutes at 4°C. Supernatants (serum) were collected and kept at -80°C until further analysis.

### Spike Protein (SP) binding antibody

His-tagged full-length SARS-CoV-2 SP (Cat. # 40589-V08B1, Sino Biological) was used as capture antigen on ELISA plates. Plates were incubated with obtained mouse serum samples for 2 h, followed by detection using HRP-conjugated goat anti-mouse IgG antibody (Cat# 05-4220, Invitrogen). Absorbance was measured at 450 nm by a Multiskan FC Microplate Photometer (Fisher Scientific) in an ELISA plate reader.

### Statistical analysis

Protection categories were defined to demonstrate the substantial heterogeneity among hmAb treatments without focusing on individual hmAbs, based on the percent-survival of mice treated with each antibody at the 1.5 mg dose (high: ≥80% survival, moderate: <80% and ≥50% survival, and low <50% survival. Survival curves were estimated using the Kaplan-Meier estimator with Greenwood’s formula for the 95% confidence intervals.

Principal component analysis (PCA) was conducted using mouse weights in grams on their original scale, recorded each day; weight after euthanasia was imputed as the final living weight for the purposes of PCA. Ellipses represent the 95% CI on the center of each group’s PC1 and PC2 coordinates.

Antibody neutralization and affinity were analyzed following a log transformation. For single biomarkers, 5-fold cross-validated receiver operator curves (ROC) were generated by computing sensitivity and specificity for prediction of survival, at every level of the marker. For multivariate biomarker combinations logistic regression was used to predict mouse survival. For group-level variables (e.g., *in vitro* neutralization) this required that the prediction for each mouse in a group was the same, since there were no mouse-level measures available. The 95% CI on the area under the cross-validated ROC curve (CV-AUC) was estimated using the non-parametric Delong’s method.

## RESULTS

### SARS-CoV-2 infection association with morbidity and mortality in K18 hACE2 transgenic mice

As part of the effort to rapidly and accurately identify hmAbs effective against SARS-CoV-2, we designed an experiment using K18 hACE2 transgenic mice (**Fig. 1A**). In this study, 206 hmAbs from the CoVIC-DB repository, known to investigators only by their COVIC identification number, were evaluated in a SARS-CoV-2 challenge model. Daily weights and observations were collected throughout the duration of the study (10 days).

**Figure 1.**
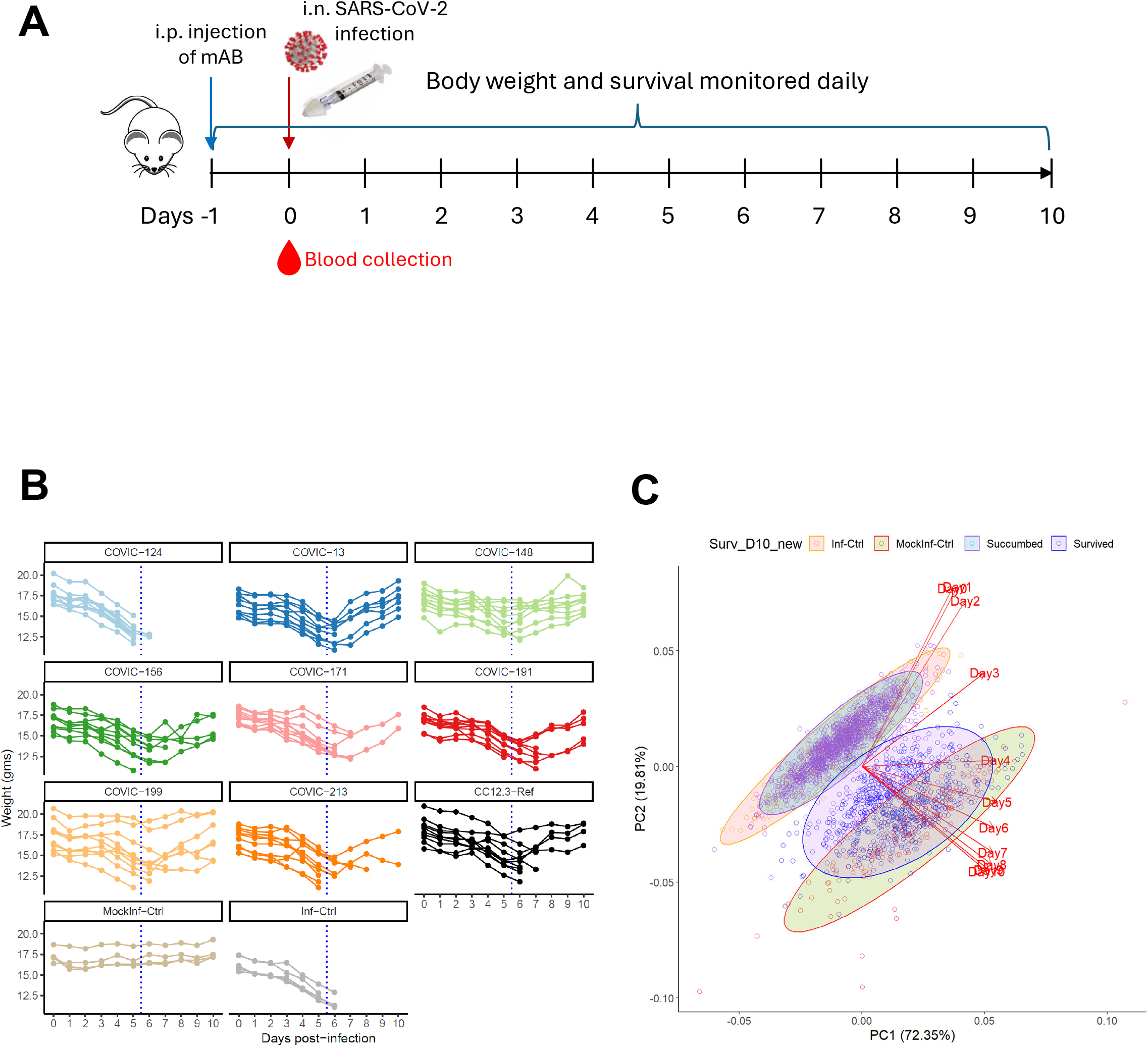
Anti-SP (hmAbs) improve survival and weight change in K18 hACE2 transgenic mice. **(A)** Study design. Mice were separated into groups of 10 and treated on day -1 by intraperitoneal (i.p.) injection with candidate hmAbs submitted to the CoVIC-DB. Intranasal (i.n.) challenge with a lethal dose (10^5^ PFU/mouse) of SARS-CoV-2 WA-1/US took place on day 0. Mice were weighed daily following challenge and monitored for symptoms. Serum samples were collected immediately prior to challenge, as indicated by red blood drop. **(B)** Mouse weight trajectory (in grams) over the 10-day experiment, shown for a set of representative hmAbs and controls. MockInf-Ctrl refers to mice mock-infected with PBS; Inf-Ctrl refers to untreated mice (mock-treated with PBS); CC12.3-Ref refers to a previously tested hmAb used as a reference control. **(C)** PCA results are shown for weight trajectories over the 10-day experiment. Circles represent the PCA score for each individual mouse and are colored based on respective group: untreated infection-control mice in yellow; untreated mock-infection control mice in red; surviving treated mice in purple; and non-surviving treated mice in blue. Shaded ellipses represent the distribution in results for each group within a 95% confidence interval, with infected control mice in peach, mock-infected mice in green, surviving mice in purple, and mice that succumbed to infection in blue. Eigenvectors, shown here as red lines, represent the changing relationship between mouse body weight and survival outcome over time. Missing body weights from mice that succumbed to infection were assigned their terminal weight.

All untreated control infected mice rapidly lost weight following challenge and succumbed by day 6 post-infection, while untreated mock-infected mice all survived and showed little change in weight over 10 days (**Fig. 1B**). A range of intermediate phenotypes were observed in mice following treatment with the different hmAbs, including slower decline in body weight, weight loss recovery, and extended survival. Results illustrated broad heterogeneity in effectiveness of different hmAb treatments, with many demonstrating improved outcomes over the reference hmAb control CC12.3.

To further explore the different weight trajectories associated with infection, hmAb treatment, and survival, we used principal component analysis (PCA) of daily mouse weights (**Fig. 1C**). The first two principal components explained 72% and 20% of the variance, respectively, suggesting that most of the variance over 10 days could be described adequately by two dimensions. Mice in the infection control and the mock-infection control groups were easily distinguished in the PCA embedding, allowing us to establish an axis of severity; mice in the hmAb treated groups tended to fall along this axis and between the two control groups. Among mice that died, there was little difference in weight loss between those that were and were not treated with hmAbs. In contrast, there was a large difference in weight trajectories between the hmAb-treated mice that succumbed to infection and those that survived, though with substantial variability in the latter group. The overlap between the treated surviving mice and the mock-infected control mice (which also survived) was less pronounced, suggesting that there was a gradient of disease severity within the mice that survived. Many of the treated mice resembled the mock-infected mice suggesting that at least some of the hmAbs were highly potent against SARS-CoV-2 infection. PCA loading vectors representing the mouse weights of the first few days were roughly orthogonal to the severity axis, indicating that the original baseline weight and the early weight change were not correlated with severity. By day 5 post-infection, vectors appeared to align with the axis of severity, suggesting that weight change from day 5 post-infection onward may be a predictor of outcome.

### Infusion of anti-SP hmAbs decreased morbidity and improved survival

To describe the relationship between survival and weight change, we categorized the hmAbs based on the proportion of mice protected in the associated groups, ranking them into high (≥80% survival), moderate (<80% and ≥50% survival), and low (<50% survival) protection categories (**Supplemental Data**). We then estimated survival probability over time for the mice in each of the categories (**Fig. 2A**, Kaplan-Meier estimator). Mock-infected mice exhibited 100% survival probability, while infected untreated control mice all succumbed by day 7 post-infection. Mice treated with high ranking hmAbs had a higher probability of survival than mice treated with the reference hmAb CC12.3, while mice treated with moderate and low protective hmAbs had poorer survival probability.

**Figure 2.**
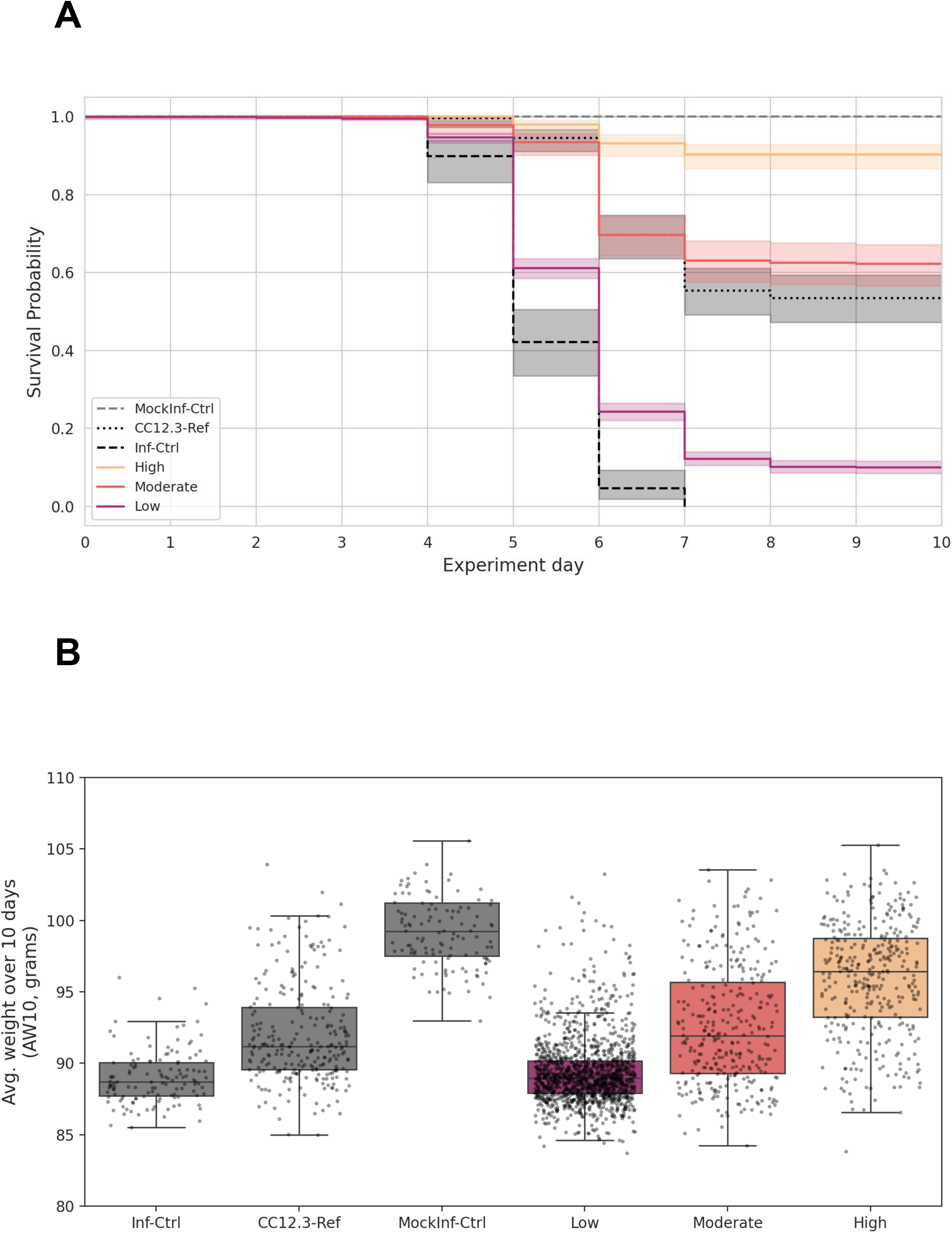
hmAbs demonstrate a broad range of protection. (**A**) Kaplan-Meier curve of survival probability (Y-axis) over time (X-axis) is shown for mice in control groups, reference hmAb CC12.3, and protection categories of hmAbs (high, n=35: ≥80% survival, moderate, n=32: <80% and ≥50% survival, and low, n=139: <50% survival). Colors distinguish among groups, with shading representing the respective 95% confidence interval. **(B)** Average weight over ten days (AW10) relative to baseline weight is shown for each group from left to right in order of decreasing effectiveness. Dots represent individual mice. Box plots show the median and interquartile range calculated for each COVIC protection category and control group.

We then computed average weight change over ten days (AW10) for each mouse relative to its baseline weight (**Fig. 2B**). Higher AW10 reflected less morbidity during the experiments, with a score of 100 representing a stable weight trajectory from baseline to day 10 post-infection. After mock-infected untreated control mice, mice treated with high protection hmAbs had the highest median AW10 scores, followed by mice treated with the reference hmAb CC12.3. Mice treated with moderate and low protection hmAbs showed little difference in morbidity trends from the infected, untreated control mice.

We additionally stratified AW10 by mouse survival (**Fig. S1**). For mice that succumbed to infection, the AW10 was similar across all three protection categories; however, for surviving mice the AW10 was higher among hmAbs that provided greater protection. For mice treated with hmAbs ranked in the high protection category, their AW10 score approached that of the mock-infected untreated mice. Together these analyses indicated that weight changes following treatment differ by hmAb treatment group, even among mice that survived, suggesting that weight change in addition to survival is an important measure of hmAb protection.

### Survival was associated with higher serum levels of anti-SP hmAb on the day of challenge

We hypothesized that the serum levels of hmAb binding to the SARS-CoV-2 SP would be associated with outcome. To test this, we used an anti-SP IgG ELISA to measure SP binding from mouse serum samples collected 24 h post-treatment, prior to SARS-CoV-2 challenge. Measuring SP binding levels from each individual mouse also provided an opportunity to assess mouse-level variability in the challenge model.

We first investigated associations at the group-level, comparing the geometric mean anti-SP binding level for each hmAb tested, categorized by treatment protection category. We observed here that SP mean binding level increased for each protection category from low to high (**Fig. 3A**). We also looked at the SP binding levels for each individual mouse to assess correlations with mouse survival (**Fig. 3B**). Median hmAb binding strength for each individual mouse was stronger for surviving *vs*. non-surviving mice within the same protection category. However, among the non-surviving mice, we saw that the median value for those treated with moderate hmAbs was higher than the median value of surviving mice treated with low protection hmAbs. Furthermore, many mice that succumbed to infection had high levels of SP binding hmAbs and conversely, there were also many surviving mice with low SP binding levels. These findings may be partially explained by technical variations or biological variability within the mice; however, these also suggest that anti-SP hmAb binding alone may not be sufficient at predicting mouse survival.

**Figure 3.**
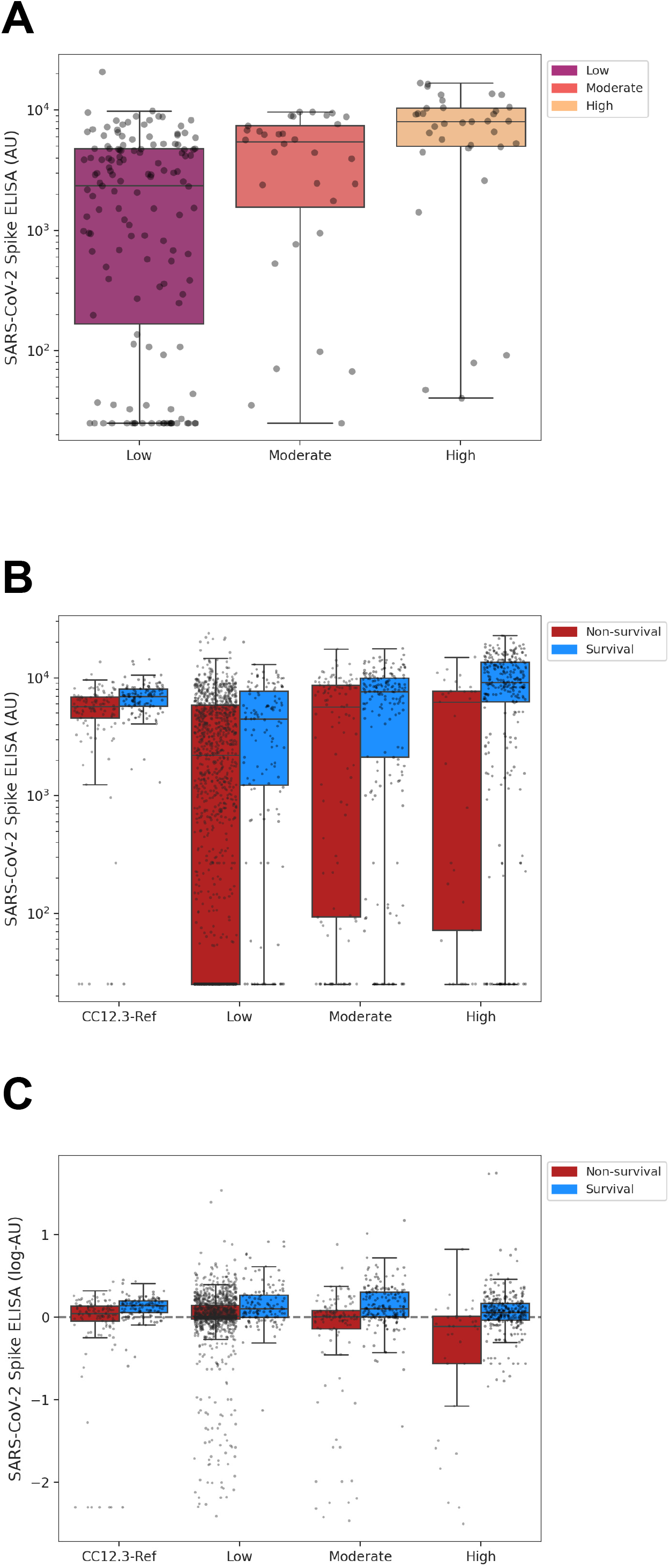
Survival is associated with anti-SP binding levels measured on day of challenge. ELISA was used on serum samples taken at time of challenge to evaluate hmAb binding strength to SP. **(A)** Geometric mean SP binding values were calculated for each treatment group and plotted by protection category (high, n=35; moderate, n=32; low, n=139). Each dot represents the mean SP binding level for a specific hmAb treatment group (n=10); colors reflect assigned protection category. **(B)** Challenge-day SP binding hmAb levels for each individual mouse. ELISA values for binding to SP on day of challenge are shown on Y-axis. Each circle represents one mouse, with treatment categories shown on the X-axis and separated into survival (blue) *vs*. non-survival (red). Boxplots reflect median and interquartile range. **(C)** Residual challenge-day SP binding hmAb level for each mouse is shown after subtracting the respective treatment group mean calculated in B (subtraction on log-scale). Residual ELISA levels (Y-axis) shown here in log_10_ units.

To differentiate between mouse-level variability and that induced by different hmAb treatments, we calculated each mouse’s residual hmAb binding level by subtracting the respective group mean from the individual binding values (**Fig. 3C**). We observed that within each protection category, the median residual binding levels were higher for mice that survived than for those that succumbed to the infection, and that across all defined treatment categories, there were mice with high levels of hmAbs that nonetheless succumbed to the infection.

### Survival can be partially predicted by the neutralization potency and SP binding level measured *in vitro*

Despite the technical and biological variations that we observed among individual mice, the correlation between SP mean and protection category suggested that average hmAb binding strength could be predictive of treatment outcome. To evaluate the strength of the relationship, we computed receiver operating characteristic (ROC) curves for SP mean as a classifier for prediction of survival (**Fig. 4A**). As part of this analysis, we also included results from the *in vitro* assays that were performed on the hmAbs as part of their submission to the CoVIC-DB. This allowed us to make head-to-head comparisons among various types of data to assess how each may contribute to survival.

**Figure 4.**
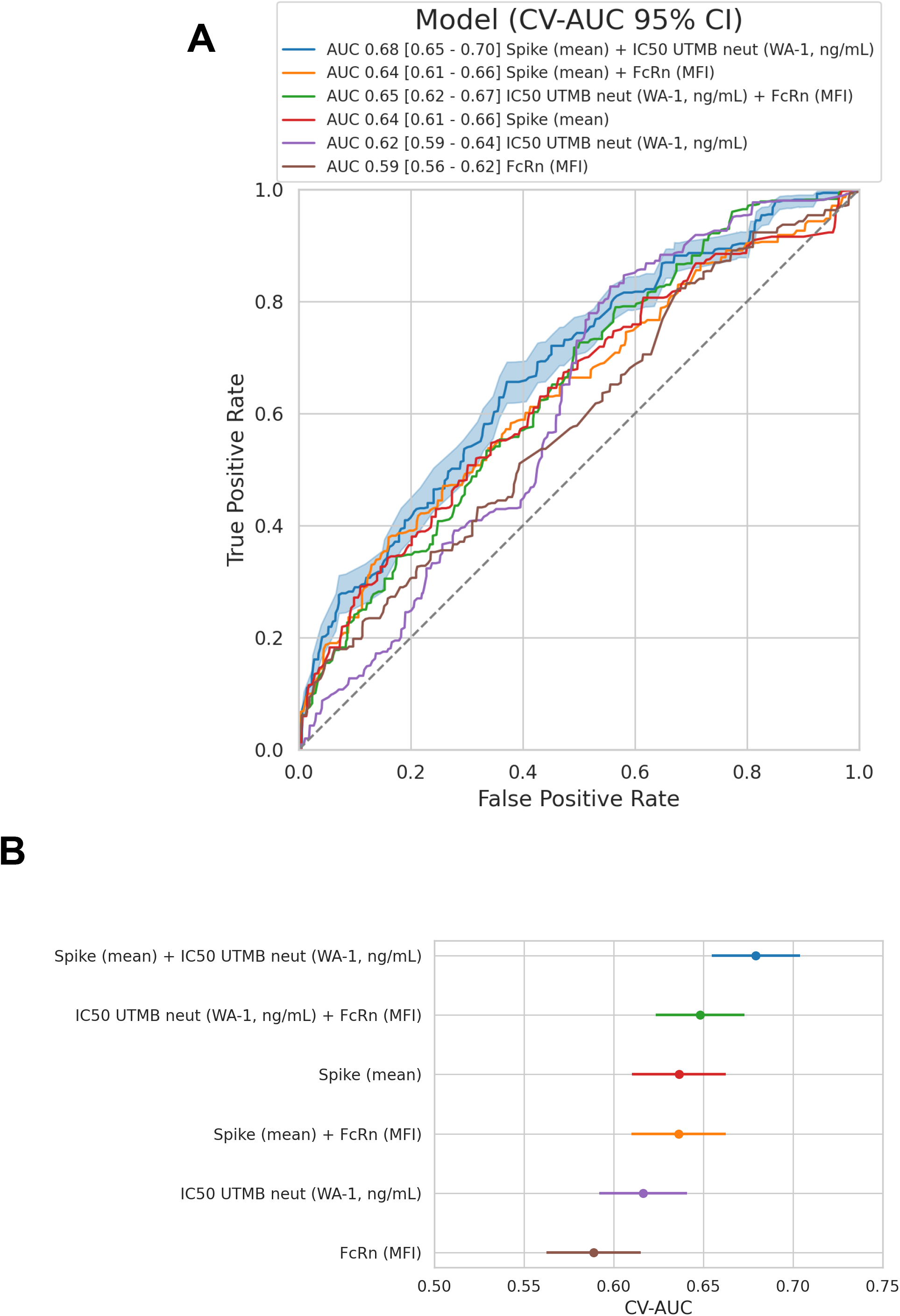
Survival is only partially predicted by *in vitro* measurements. *In vitro* data collected from the CoVIC-DB for each hmAb were evaluated for how well they predict survival in comparison to challenge-day mean SP ELISA data collected here (“Spike (mean)”). **(A)** A cross-validated ROC curve is plotted for each variable or variable set, indicating ability to classify mouse survival. Cross-validation was conducted by leaving one treatment group out in each iteration. A 95% confidence interval (CI) is shown for the model with the highest area under the curve (AUC). All AUCs are provided in the legend with 95% CI. **(B)** Forest Plot of CV-AUC scores with 95% CI. IC50 UTMB neut WA-1, concentration of hmAb needed to neutralize 50% of infection by authentic wild-type SARS-CoV-2 D614 strain WA-1/US measured at the University of Texas Medical Branch (UTMB); FcRn (MFI), mean fluorescence intensity of binding to neonatal Fc Receptor.

Cross validated ROC curves showed that even though the SP mean variable (group-level mean SP binding) was the best individual predictor, its Area Under the Curve (AUC) was quite low at 0.64 [0.61 - 0.66]. Prediction was only modestly improved when combined with data from viral neutralization assays (using multivariate logistic regression), increasing to 0.68 [0.65 - 0.70]. When SP mean data were combined with the residual binding data (**Fig. S2**), AUC also increased slightly to 0.67 [0.65 - 0.70]. This suggests that the residual SP binding data are at least as useful as the information from the neutralization assays. Consistent with previous observations, residual SP binding data alone were not predictive of outcome, but the increased AUC score indicated that mouse-level variability contributed somewhat, albeit modestly, to improving prediction. All *in vitro* assay readouts, either alone or in combination, yielded low to moderate AUC scores, showing that survival was only partially predicted with these assessments.

### Integration of *in vivo* and *in vitro* metrics allowed for ranking of highest performative hmAbs

The *in vitro* data in the CoVIC-DB were combined with the *in vivo* mouse protection data to compare the effectiveness of the hmAb candidates. Focusing first on the 35 hmAbs that protected at least 80% of the mice in their treatment groups, we generated a heatmap to display them based on performance across all results (**Fig. 5**). Most of these protective hmAbs had high scores in SP binding and viral neutralization assays. However, this was not always the case, e.g., hmAb COVIC-78 had poor *in vitro* neutralization and SP binding yet yielded 100% protection. Other hmAbs with promising *in vitro* data failed to protect *in vivo* (**Fig. S3**).

**Figure 5.**
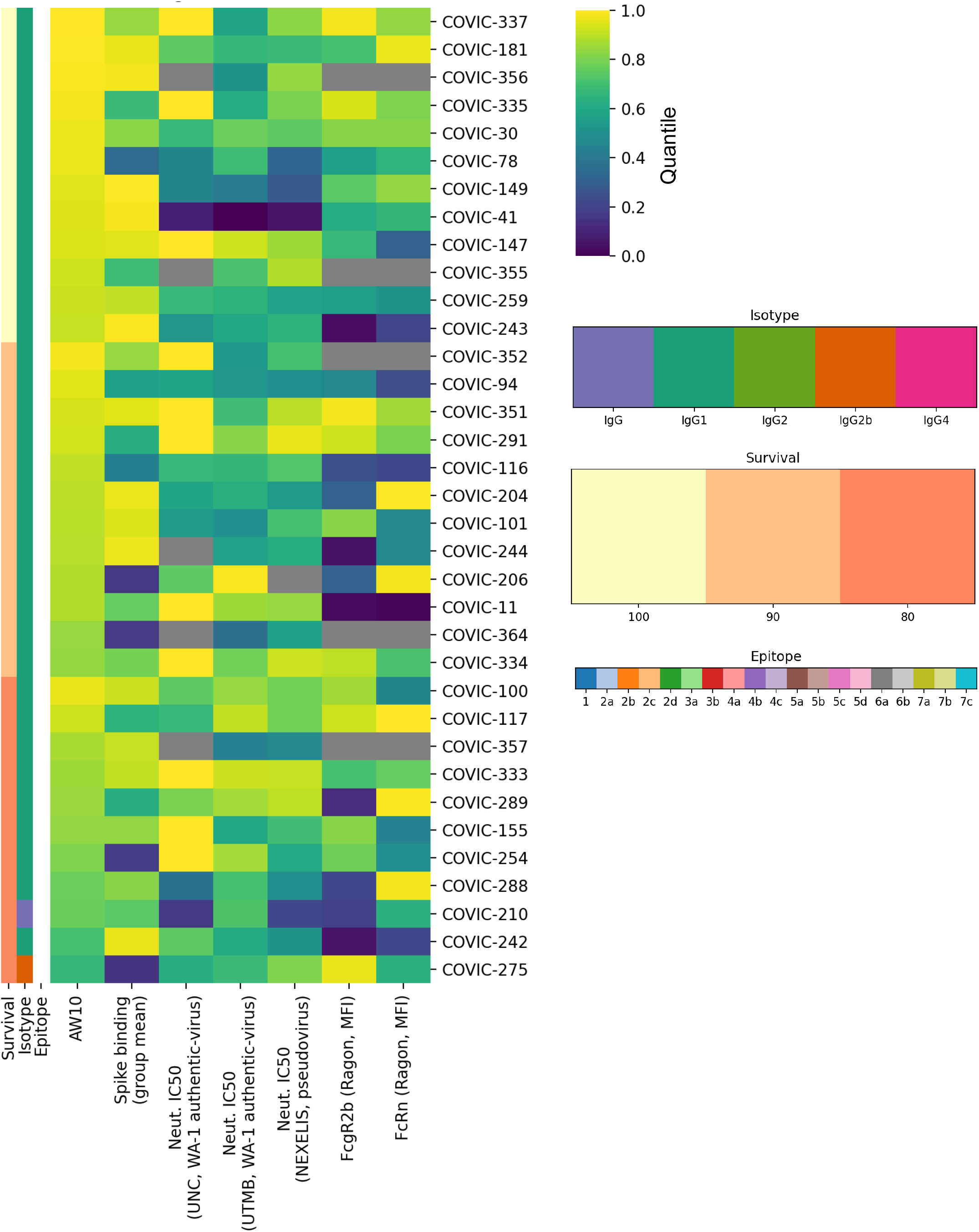
Heat map for hmAbs with greater than 80% protection. The hmAbs that protected at least 80% of mice are listed by row in decreasing order of protection. Columns represent qualifiers (e.g., mouse survival status) and scores for assay results. Color scores are shown as a percentile on a scale from 0 (indigo) to 1.0 (yellow). Legends for percent mouse survival, isotype, and epitope (if known) are shown above. AW10, percent mouse body weight change over 10 days; RBD (ELU), ELISA units for binding to SP Receptor Binding Domain; IC50 UTMB, concentration of hmAb needed to neutralize 50% of infection by authentic wild-type SARS-CoV-2 D614 strain WA-1/US measured at the UTMB; IC_50_-Pseudovirus, concentration of hmAb needed to neutralize 50% of infection by pseudovirus; FcgR2b (MFI), mean fluorescence intensity of binding to Fcγ receptor 2b; FcRn (MFI), mean fluorescence intensity of binding to neonatal Fc Receptor.

## DISCUSSION

Outbreaks of deadly coronaviruses such as SARS-CoV, MERS-CoV, and SARS-CoV-2 in recent decades remind us of the urgency for effective therapeutics in the face of rapidly changing global health crises, especially in resource limited settings. Here, using the K18 hACE2 transgenic mouse model, we screened a panel of hmAb candidates from the CoVIC-DB and identified many that were highly protective against a lethal SARS-CoV-2 challenge. Notably, protection *in vivo* did not necessarily correlate with *in vitro* assay measurements. Several highly protective hmAb candidates – some conferring 100% protection *in vivo* – were unremarkable by at least one (and sometimes all) of the *in vitro* experiments. Furthermore, even though survival correlated with mean anti-SP binding level, our ROC analysis showed that like other *in vitro* measurements, anti-SP binding was a poor predictor of protection, even when modestly improved upon by incorporating data from the other experiments. Together the findings demonstrate that the *in vivo* model provides distinct information about hmAb candidates.

To evaluate how the mouse model performs to predict hmAb efficacy against SARS-CoV-2 infection, we also assessed mouse-level variability, which would drive the sample size required to test other potential hmAbs in future studies. We observed mice in high-protective treatment groups that succumbed, with several having lower-than-average anti-SP binding at challenge day (*i*.*e*., residual SP binding); this could be due to variability in hmAb dosing or in hmAb clearance by the host. However, overall residual binding SP levels were not predictive of survival in the ROC analysis, suggesting that the mouse-level variability was low when compared to the treatment-level hmAb effects. Mouse-level variability in weight trajectories seemed to carry valuable information in addition to survival alone. This was evident from the PCA analysis, which showed a gradient of mouse weight trajectories along a severity axis, and the observation that AW10 scores were higher in the high-protective *vs*. moderate-protective category, even among mice that survived. This showed that mouse weight trajectories could serve as an early outcome of the *in vivo* model and provide a more granular readout of the hmAb effectiveness when compared to survival alone.

While the mouse model was able to identify protective hmAbs that would not have otherwise been detected by *in vitro* screening experiments, there were some limitations that warrant mention. First, our data only reflect hmAb prophylactic treatment at a single dose given to female mice. Pilot studies using the reference CC12.3 hmAb informed the design for the trial described here, but it would be useful to assess effectiveness of the hmAbs under varying concentrations, and/or alternate dosing schedules, including administration after SARS-CoV-2 challenge. Future experiments could also incorporate challenge by different viral variants or increased viral titers and include both male and female mice to account for the impacts of sex on hmAb treatment outcomes. Another limitation was that the hmAb structure and amino-acid sequence were blinded as part of their inclusion in the CoVIC-DB, a criterium set up to incentivize submissions from parties interested in protecting their intellectual property. Though for many of the hmAbs, isotype and epitope targets were known and discussed in Schendel et al. (23), the hmAb blinding made it difficult to further investigate the results, particularly when the *in vitro* measures and *in vivo* protection were discordant.

Transgenic expression of the hACE2 receptor in airway cells in the K18 hACE2 mouse model allowed us to study hmAb candidates in a mouse with infection pathogenesis that mimics COVID-19 in humans (18). The ACE2 receptor is used as a point of entry by a variety of alpha and beta coronaviruses, many of which reside in wild animal populations and can recognize ACE2 variants from more than one host species (24–27). As humans continue to expand their interaction with the natural environment, zoonotic spillover events with pandemic potential will remain an issue (28, 29). The ability to impart swift and accurate screening of therapeutic treatments in the face of such disease outbreaks will be crucial. While *in vitro* experiments provide important characteristics of hmAb candidates, here we have shown that the K18 hACE2 mouse model can meaningfully differentiate protective hmAbs when given as a prophylactic treatment. This model provides a unique evaluation of potential hmAb performance and should be considered as a valuable complement to *in vitro* screening of therapeutic candidates.

## AUTHOR CONTRIBUTIONS

BB, PAP, AH, JIG, AG-V, CY, J-GP, BM and OR performed experiments and data analyses. TS, DH, SLS, EOS, LMS, JBT and AF-G performed experimental design, data analyses, interpretation and/or wrote or revised the draft. SS, EOS, LMS, JBT and AF-G obtained funds for this study. All co-authors agreed to the final version of this manuscript.

## DATA AVAILABILITY

All data supporting this manuscript are included in the Supplement.

## ACKNOWLEDGEMENTS

We thank Dr. Jacqueline Kirchner at the Gates Foundation for her support and guidance during the performance of these studies. We thank all researchers and administrators involved with CoVIC that made this study possible. We thank the Texas Biomed BSL3/ABSL3 Operations Program for their commitment and support. We also thank all third parties involved in this study for facilitating the performance of research during the COVID-19 pandemic.

